# Extensive and diverse lanthanide-dependent metabolism in the ocean

**DOI:** 10.1101/2023.07.25.550467

**Authors:** Marcos Y. Voutsinos, Jillian F. Banfield, Harry-Luke O. McClelland

## Abstract

Lanthanide (Ln)-dependent enzymes have evolved roles in organic carbon metabolism despite low Ln availability in natural environments^1–8^. The oceans are the major reservoir of dissolved organic carbon (DOC) on the planet, yet the prevalence and diversity of Ln-dependent enzymes in the ocean, and their biogeochemical importance in the ocean carbon cycle is unknown. Here, we analyzed a global ocean metagenomic/metatranscriptomic dataset and found Ln-dependent methanol-, ethanol- and putative sorbose- and glucose-dehydrogenases in all metagenomes and 20% of all resolved microbial genomes, with several individual organisms hosting dozens of unique Ln-dependent genes. We find that biological methanol oxidation in the ocean is overwhelmingly Ln-dependent, and that methanol dehydrogenases are the most highly expressed Ln-dependent genes in most ocean regions, particularly in surface oceans. As Ln availability is a function of phosphate concentration and pH, Ln-dependent metabolism likely underpins complex biogeochemical feedbacks determining the efficiency of organic matter remineralization, thus impacting the oceanic DOC pool and Earth’s climate system. The widespread biological utility of Ln also explains their nutrient-like vertical concentration profiles observed in ocean waters, and shows that the preferential utilization of light lanthanides by biology must be considered when interpreting patterns of relative Ln concentrations in seawater.

## Introduction

The lanthanide elements (Ln), also known as the rare earth elements (REE), were long thought to be entirely biologically inert. This view changed in 2011 when a methanol dehydrogenase (MDH) was discovered that was upregulated in the presence of Ln^1,2^ and used Ln as a cofactor^3^, ultimately leading to the discovery of an obligately Ln utilizing bacterium^4^. These unexpected findings spurred a decade of intense research into Ln-dependent enzymes. So far, a handful of Ln-dependent methanol and ethanol dehydrogenases have been characterized, all of which use the lighter of the Ln series (mostly La, Ce, Pr, and Nd), and belong to the group of dehydrogenases (DHs) that employ the redox cofactor pyrroloquinoline quinone (PQQ) in the active site. The utility of Ln within the cofactor complex stems from their electrophilicity, replacing the function of Ca, on which all PQQ-DHs were previously thought to depend^9^. Although of great scientific and industrial interest, at first Ln-PQQ-DHs were assumed to be biogeochemically unimportant given the billion-fold greater concentrations of Ca over Ln in natural environments. Yet there has been growing evidence that Ln-PQQ-DHs are superior to their Ca-dependent counterparts^10^, and that Ln-dependent metabolism may thrive in certain natural settings^5–8,11,12^. However, the global distribution and bio(geo)chemical importance of lanthanide-dependent metabolism in nature remains virtually unknown.

In this work, using a publicly available global metagenomic and metatranscriptomic database, we explore the extent, distribution and diversity of Ln-dependent metabolism in the surface ocean. We find that Ln-dependent enzymes are ubiquitous and are both more highly expressed and far more diverse than previously thought. We show that non-methanotrophic methanol dehydrogenase (MDH) is the most highly expressed Ln-dependent enzyme in the surface ocean, hosted by members of an unknown alphaproteobacterial clade closely related to the order Rickettsiales. Ln-dependent MDHs are by far the most highly expressed methanol oxidation genes in the ocean, highlighting the central role that Ln plays in the marine C1 carbon cycle. At greater water depths, Ln-dependent metabolism mostly involves longer chain simple alcohols. We also identify a handful of organisms as likely lanthanophore producers, which may be important biotechnology targets for lanthanide biomining and purification. Furthermore, we find that the spatial distribution of Ln-dependent metabolism is strongly related to phosphate concentrations and pH in the modern surface ocean, which may be due to a combination of the limiting effect of PO_4_^3-^ on the concentration of Ln in seawater ^13^, enhanced scavenging efficiency at low pH ^14^, and greater depletion of Ln in regions of high productivity via biological uptake and export. This ubiquitous biological role for Ln introduces a mechanistic coupling between the cycling of lanthanides in the ocean and the marine carbon cycle: bacterial uptake is known to partition Ln with preferential removal of the lightest of the series; while the availability of Ln regulates an organic carbon remineralization flux that may constitute a significant leak of CO_2_ from the oceanic dissolved organic carbon pool.

### PQQ-DHs in the ocean are primarily Ln dependent

We used a hidden Markov model (HMM) to detect PQQ-binding dehydrogenase enzymes (PQQ-DH) in the TARA Oceans Ocean Microbial Reference Gene Catalog version 2 (OM-RGCv2)^15^. This global dataset includes metagenomic and metatranscriptomic sequence data for samples from the surface ocean (SO) and the deep chlorophyll maximum (DCM) in the epipelagic zone, and from the mesopelagic zone (MES). The catalytic binding sites of lanthanide-dependent PQQ-DH enzymes contain an additional aspartate residue absent from calcium-dependent homologs^16^. Recent studies have confirmed that the aspartate residue is essential for lanthanide coordination and activity in PQQ-ADH Type 1^16^ and PQQ-ADH Type 2a and 2b^17,18^ enzymes, and that Ln is essential for these enzymes to function. The presence of this sequence motif is therefore taken to be a direct indication of lanthanide binding^10,16^ (Methods; Fig. S1).

We identified 6,886 PQQ-DH proteins in the microbial metagenomes (Fig. 1A), of which 56% contained the lanthanide binding motif, 11% contained the homologous calcium binding motif, and 24% contained a homologous sequence to the Ln and Ca binding domains but with neither the diagnostic Ln or Ca binding residues. The remaining sequences were missing at least one amino acid from the catalytic cofactor binding domain. This latter group of sequences are included in the protein trees (Fig. 1A), but as the metal cofactor could not be determined, they were excluded from further analyzes. Sequences were classified to protein family level according to InterPro^19^, resulting in 17 PQQ-DH families, 14 of which were primarily or entirely Ln dependent (Fig. 1C).

**Figure 1.**
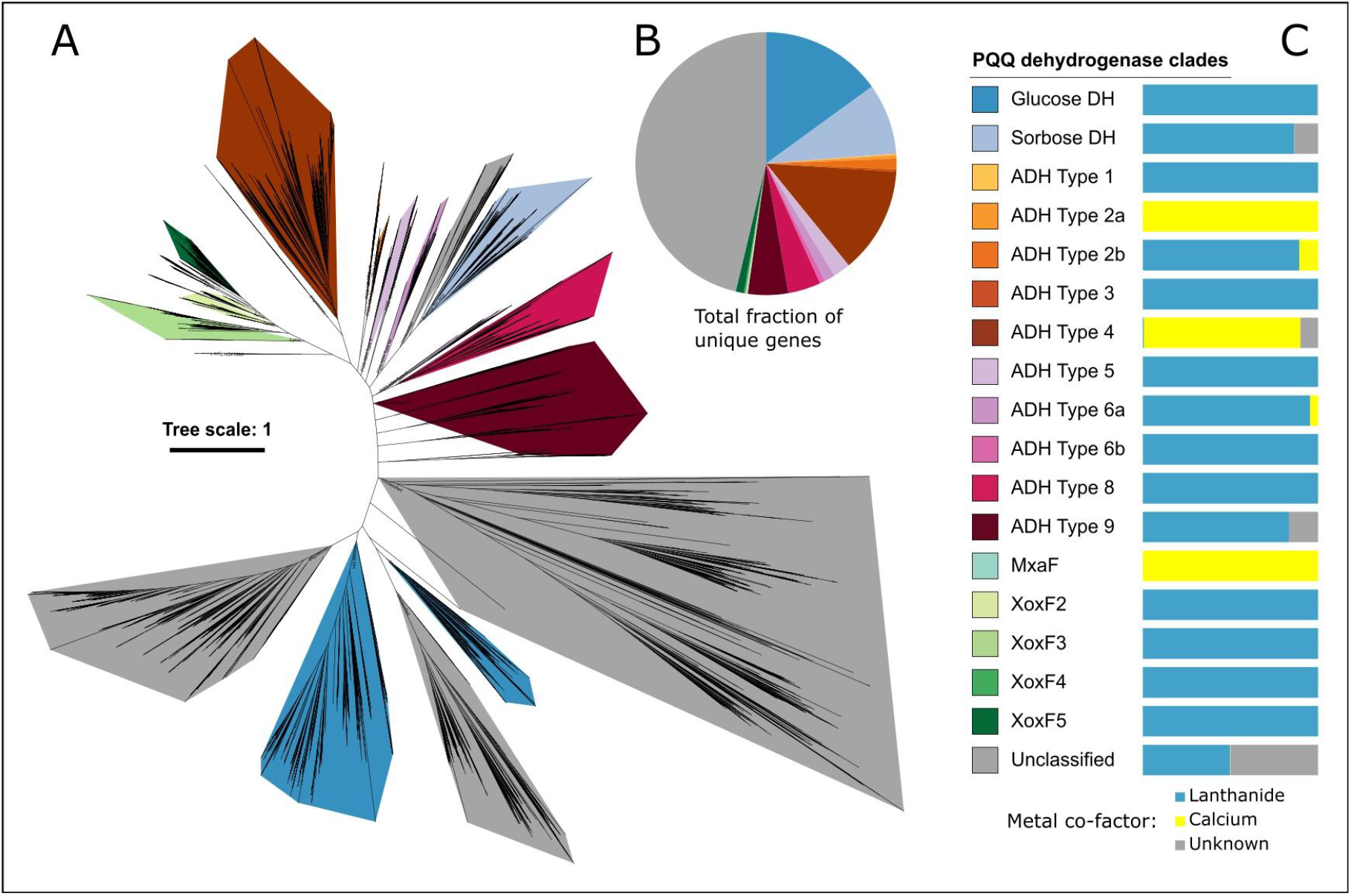
Analysis of 6886 dereplicated PQQ dehydrogenase genes from Tara Ocean metagenomes and metagenome assembled genomes (MAGs). **(A)** Unrooted tree representing manual phylogenetic classification of PQQ dehydrogenase clades across SO, DCM and MES metagenomes. **(B)** Fraction of all unique PQQ dehydrogenases represented in each clade. **(C)** Legend for A and B, and fraction of sequences in each clade containing lanthanide, calcium or unknown binding domains.

Spatial variation in the relative abundance of PQQ-DH genes in the metagenomes is modest (Fig. S3 and Fig. S4). However, community level expression is far more variable, falling into 3 distinct regimes: 1. A high temperature surface ocean regime dominated by the XoxF5 MDH; 2. A low temperature surface ocean regime dominated by the XoxF4 MDH and ADH type 3; and 3. A deep water regime, dominated by ADHs types 6a, 2a and 2b (Fig. S3 and Fig. S4). The samples from the DCM occupy the transitional zone between these regimes. Ln-dependent metabolism in the surface ocean is dominantly methylotrophic, whereas oxidation of longer chain simple alcohols (ethanol, propanol or butanol) appears to dominate at depth (Fig. S5). Deep samples express a higher fraction of unclassified sequences whose substrates are unknown.

We also explored the Bacterial and Archaeal metagenome assembled genomes (BacArcMAGs; MAGs) dataset^20^, which is based on TARA oceans metagenomes from the SO and DCM. We found that 14 / 33 phyla and 20% of all MAGs contained Ln-PQQ-DHs. Three Acidobacteria genomes and a group of fifteen Pseudomonadales UBA9145 genomes contained an unusually large repertoire of Ln-PQQ-DHs, with up to 70 unique Ln-dependent genes in a single genome (Fig. 2).

**Figure 2.**
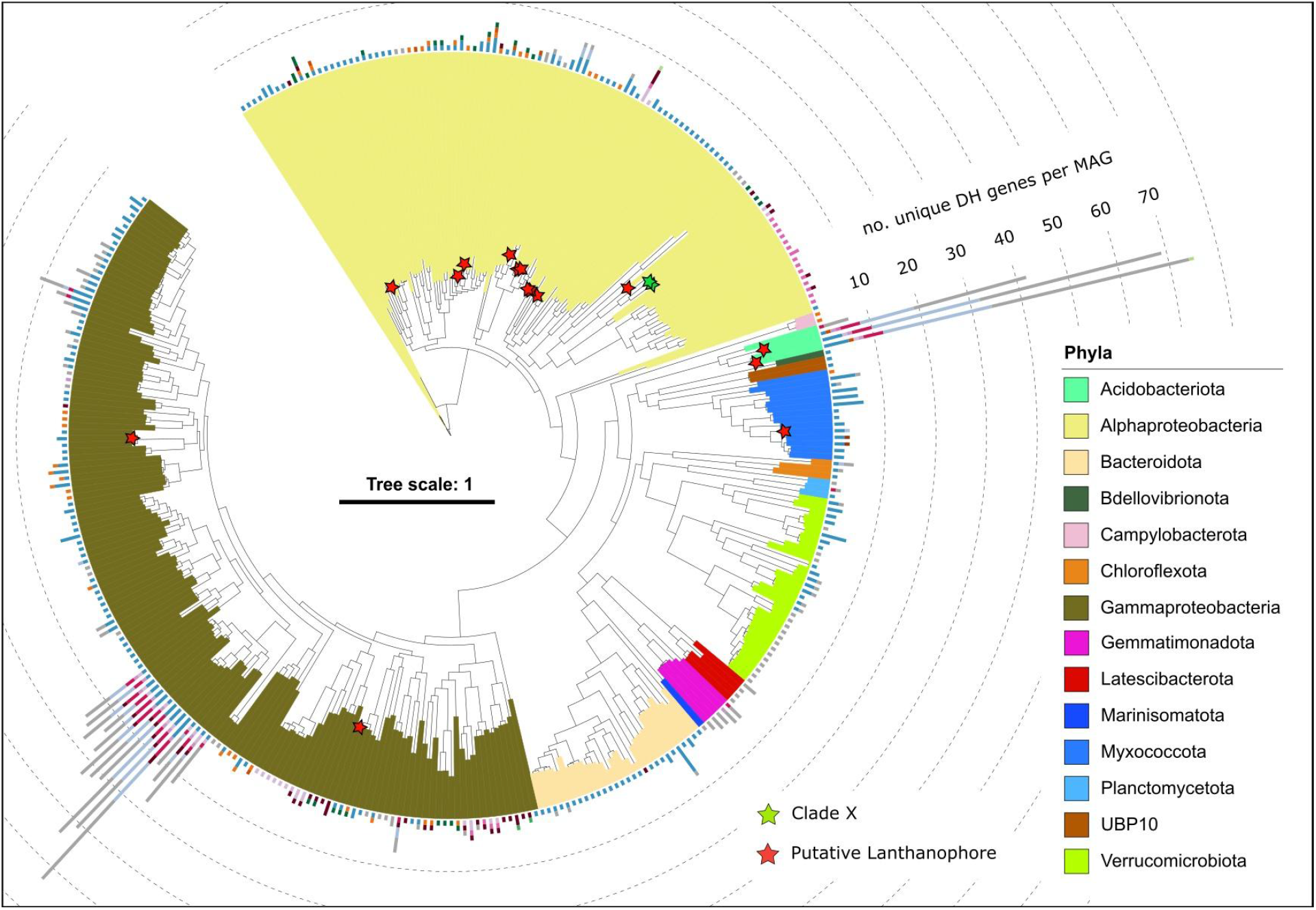
Phylogenetic tree of a concatenated alignment of 16 ribosomal proteins from MAGs from SO and DCM containing Ln-binding PQQ-DH, with number of unique PQQ-DH genes for each MAG. Red stars are placed beside MAGs containing putative lanthanophores. MAGs without Ln-binding PQQ DHs are omitted. A green star is placed beside Clade X. The MAG clade X contains the most highly expressed Ln-binding PQQ DH gene, see Fig. S6 & S7. Note: colors used in the tree correspond to taxonomic classification and bars around the edge correspond to PQQ-DH clades – legend in Fig. 1C. Tree scales represent the length of branch in which there is a 10% difference between two sequences.

Ca concentrations in seawater are around 10^-2^ M ^21^ while Ln concentrations are on the order of 10^-11^ M ^22^, yet the vast majority of PQQ-DH enzymes we detected are Ln-dependent (Fig. S2).

This result, while surprising, supports several previous findings that PQQ alcohol DHs (PQQ-ADHs) are primarily lanthanide dependent ^5–8,12,23^, but also expands this view to a greater diversity of protein families, and a more nuanced picture involving metal cofactor heterogeneity within families previously thought to bind exclusively Ca (e.g., ADH type 4) or Ln (e.g., ADH type 2b). The dominance of Ln-over Ca-dependent PQQ-DHs in nature is consistent with the experimentally demonstrated superiority and preferential use of Ln over Ca in the laboratory ^10,24^.

### Diverse Ln-dependent glucose and sorbose dehydrogenases

Glucose DHs (GDHs) and sorbose DHs (SDHs) respectively represent 13% and 8% of all PQQ-DHs in the metagenomes (Fig. 1B). Almost all of the GDHs and the vast majority of the SDHs contain the Ln-binding motif (Fig. 1C), making them the most diverse groups of Ln-PQQ-DHs in the dataset. To our knowledge this is the first time that putative Ln-dependent GDHs and SDHs have been identified, expanding the role of Lanthanides in metabolism beyond simple alcohols.

Many different Ln-PQQ sorbose DHs occur in the genomes of a relatively small number of organisms that possess these genes (mostly Acidobacteria and Gammaproteobacteria) (Fig. 2). By contrast, Ln-PQQ-GDHs occur in low numbers in each genome, and the organisms in which they occur are phylogenetically diverse (Fig. 2), and are geographically widespread (Fig. 3A). The Ln-dependence of PQQ-GDHs is consistent with recent work showing increased expression of GDH in a methylotroph when supplemented with Ln ^25^. Yet expression levels of GDHs and SDHs in the surface ocean are low, representing just 1% each of total PQQ-DH transcript abundance (Fig. 3C). These monosaccharide dehydrogenases therefore likely play an important metabolic role, but only in certain circumstances not captured in the data.

**Figure 3.**
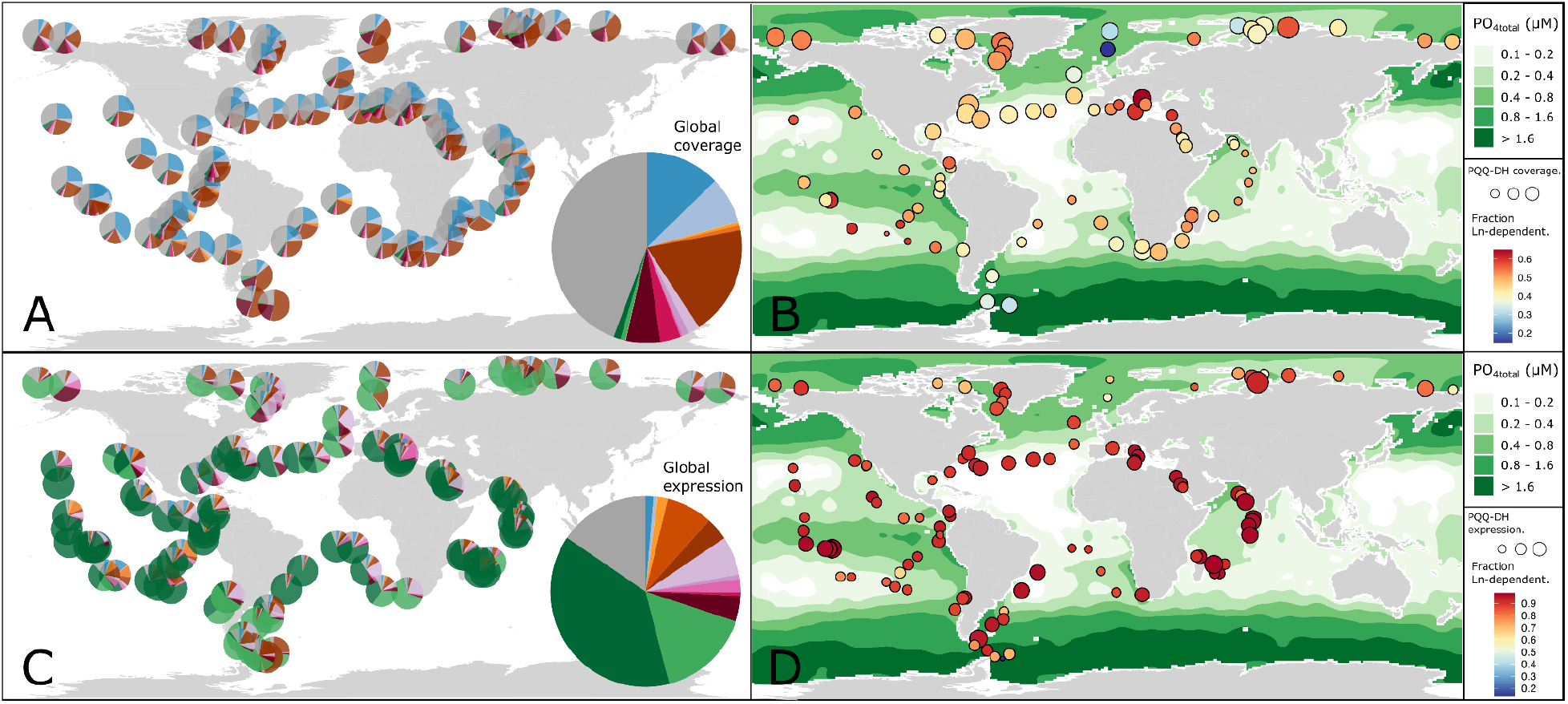
Spatial distribution of presence and expression of Ln-dependent PQQ DHs in the surface ocean. **(A)** Distribution of PQQ DH gene abundances throughout the surface ocean in community metagenomes (see Fig. 1C for color key). **(B)** Fraction of PQQ DH genes present that contain the Ln-binding motif, plotted over annual average total phosphate concentrations. **(C)** Distribution of PQQ DH transcript abundances throughout the surface ocean in community metatranscriptomes. **(D)** Fraction of PQQ DH transcripts that contain the Ln-binding motif.

### Methanol dehydrogenase dominates expression in the surface ocean

Methanol dehydrogenase (MDH) genes have low coverage in the metagenomes but dominate PQQ-DH expression, particularly in the surface ocean. Two clades, *xoxF5* (n = 63) and *xoxF4* (n = 11), which are the dominant PQQ-MDH clades reported in a variety of environments ^7,26,27^, together represent just 1% of all unique PQQ-DH genes and 1% of the gene abundance, but 55% of transcript abundance. In the surface ocean, *xoxF4* genes are more highly expressed in the high latitudes while *xoxF5* genes are more highly expressed in the low and mid latitudes (Fig. S3 and Fig. S4). The enzyme XoxF5 is kinetically superior to XoxF4, both in terms of its maximum catalytic rate and high affinity for its diverse substrates^27^. However, fast rates of methanol oxidation can be toxic if inefficient downstream processes allow an intracellular build up of formaldehyde (Fig. S5). Therefore, we hypothesize that these distinct biogeographical patterns reflect an optimized rate of methanol oxidation, with selection for the faster form, XoxF5, in warm regions where growth rates and formaldehyde removal rates are higher, and selection for the slower form, XoxF4, in cooler regions were these rates are low.

We find that XoxF4 enzymes are exclusive to the family *Methylophilaceae* of Gammaproteobacteria whereas XoxF5 enzymes are widespread across Proteobacteria, consistent with previous findings ^27^. To identify the dominant Ln-dependent organisms we created a tree of the most highly expressed *xoxF* genes from the metatranscriptomes with all *xoxF* genes identified in the MAGs (Fig. S6). The three most highly expressed *xoxF5* sequences group with *xoxF5* sequences derived from three Alphaproteobacterial genomes previously classified as Rickettsiales^20^. Further phylogenetic characterization placed these genomes in a clade with several other MAGs from the Tara oceans database also previously classified as Rickettsiales (Clade X, Fig. S7). The phylogenetic relationship between Clade X and its neighboring groups including Roseobacter, SAR116 and Rickettsiales^28^ and the Pelagibacterales (previously SAR11) were resolved with low confidence. Given their phylogenetic placement, and their appearance in the 0.22 - 3.0 μm fraction, we conclude that members of Clade X are most likely free-living, but their nature and mode of life requires further investigation.

We find that Ln and PQQ-dependent dehydrogenases are the most abundant, and by far the most highly expressed, methanol oxidation genes in the ocean metagenomes. Assuming negligible contribution from prokaryotes in the > 3 µm fraction (which are not included in OM-RGCv2), Ln-dependent MDHs comprise 80% of gene abundance and 99% of transcript abundance across all putative methanol oxidation genes in the global ocean database, including genes encoding non-Ln dependent PQQ-DHs, NAD-dependent DHs, and eukaryotic O_2_-dependent methanol oxidases (Fig. S8.) The availability of Ln is therefore central to this biogeochemically important process in the ocean.

Sources of methanol in the surface ocean are poorly understood. Ln-PQQ-MDHs are employed by some methanotrophic communities ^29,30^, however, an analysis of methanotrophy marker genes (pMMO and sMMO) in the metagenomes did not reveal any genes associated with methanol oxidation. The substrate for the Ln-PQQ-MDHs in the analyzed samples is therefore likely not methane-derived methanol (Fig. S9a,b), although this does not preclude the likely importance of methane-derived methanol in regions not captured in this dataset such as shelf regions where massive amounts of methane are released from subsurface deposits. Pectin, the primary source of methanol in the phyllosphere, which is the habitat of most well studied methylotrophs, is also in short supply in the open ocean. Single celled phytoplankton, which have been shown to produce methanol in significant amounts ^31^, appear to be the most likely source of methanol in the open ocean.

### Identification of putative lanthanophores

The hypothesized existence of lanthanophores, Ln chelators analogous to siderophores for iron ^24^, has been invoked to explain Ln utilization in aqueous environments where low Ln solubility might limit bioavailability. Lanthanophores have great potential in industry. Metallophores produced by biosynthetic gene clusters (BGC) with TonB-dependent receptors are known to play an important role in metal uptake in several species of bacteria ^32,33^. The mechanisms required for lanthanide acquisition and transport are still relatively unknown but are expected to be analogous to siderophore-mediated iron transport^34,35^. The alphaproteobacterium *Methylobacterium extorquens* AM1 has been shown to upregulate a BGC containing a TonB dependent receptor when grown with poorly soluble Nd_2_O_3_, which may be evidence of lanthanophore production^36^. We found 19 genomes containing Ln-PQQ-DHs that also contained a nonribosomal peptide synthetase BGC that contained the TonB-dependent receptor. Two of these genomes also contained over 30 lanthanide dependent enzymes, while four genomes additionally contained FecCD transmembrane protein (Type II ABC importer) and Peripla_BP_2 (substrate binding domain) in the BGC. The presence of these proteins has been shown to positively correlate with metallophore uptake^37^. Despite their significant economic value, efficient extraction and purification of Ln from raw materials remains an unsolved challenge. We highlight these lanthanophore-bearing organisms as appealing targets for use in bioleaching and biomining efforts.

### Marine biogeochemistry of the Lanthanides

Lanthanide concentrations in seawater exhibit nutrient-like vertical profiles, being low in surface waters, and increasing, and ultimately plateauing, with depth ^38^. Such profiles are typically reflective of biological uptake and remineralization at depth. The concentration profiles of Ln have been reconciled with their assumed lack of biological utility with various scavenging models. However, our findings demonstrate that Ln is indeed a nutrient, and therefore that such behavior should be expected.

To further understand biogeochemistry of Ln, we explored the relationships between environment and a) the fraction of DH genes present in the metagenomes that are Ln-dependent (fLn_G_; Fig. 4A), and b) the fraction of DH transcripts present in the metatranscriptomes that are Ln-dependent (fLn_T_; Fig. 4B). Broadly, fLn_G_ is highest in the more oligotrophic regions of the ocean, and lowest in the high-nutrient low-chlorophyll (HNLC) regions of the ocean. High values of fLn_T_ are found everywhere, but low values are only found in the HNLC regions. These ratios represent a normalization of Ln-dependent enzymes to functionally similar enzymes that use a different cofactor. The most direct explanation for differences in fLn_G_ and fLn_T_ is therefore that they are driven by Ln concentrations, which is the result of a complex balance of inputs and outputs (Fig. 5). Inputs of Ln to the surface ocean are highly spatially heterogeneous including terrigenous input from rivers ^39^, wind-blown dust, subaqueous volcanism and upwelling of nutrient-rich deep water ^40^, while outputs canonically include precipitation, and adsorption to clay minerals ^41^ and planktonic biomass ^42^, in addition to biological uptake as described here. We focus our discussion on three possible outputs of Ln from the surface ocean: 1. Biological uptake; 2. Scavenging via adsorption to sinking particles including a) phytoplankton biomass, and b) abiogenic minerals; and 3. Precipitation as LnPO_4_.

**Figure 4.**
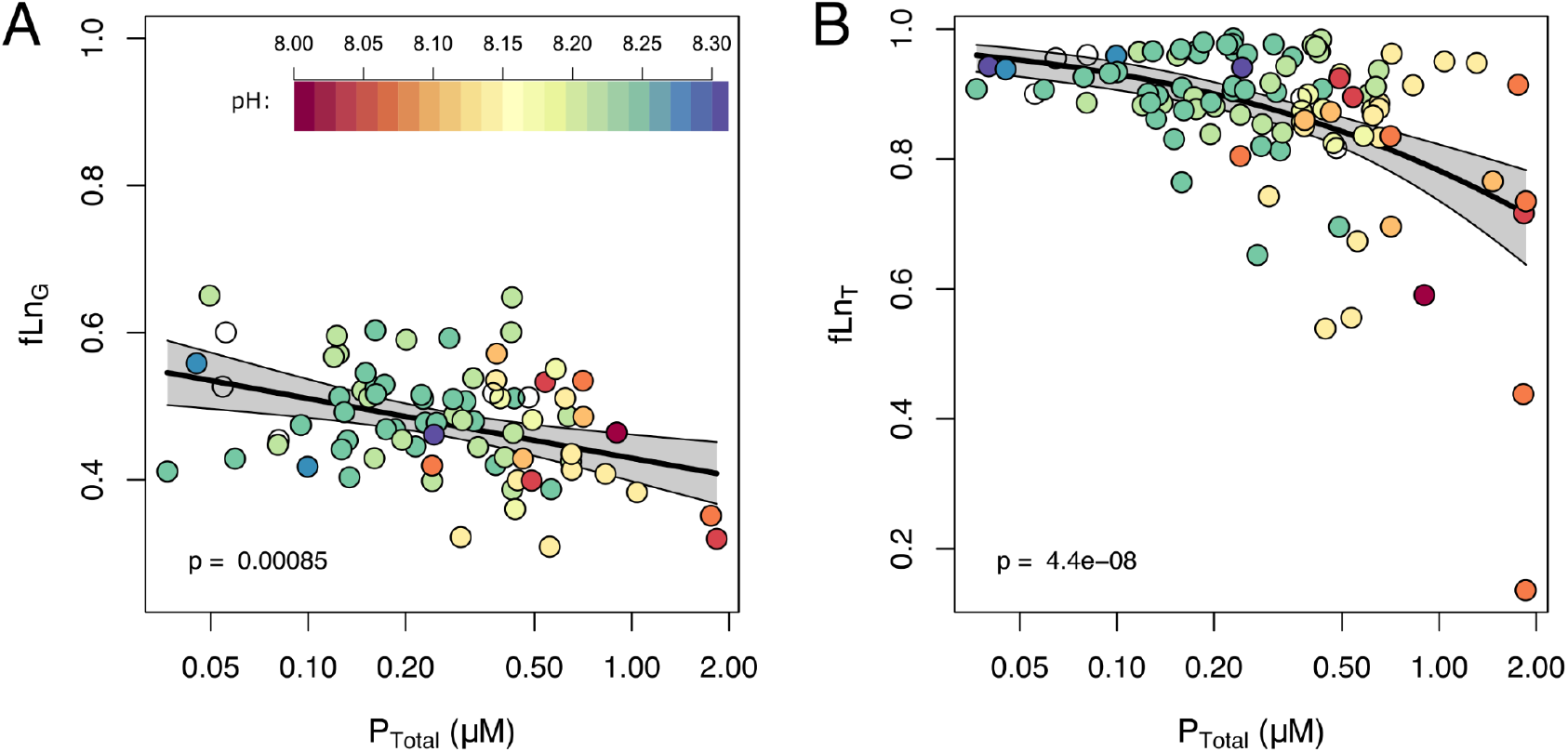
**(A)** Fraction of PQQ DH genes present that contain the Ln-binding motif (fLn_G_) plotted against the annual average total phosphate concentration at the location of sampling. **(B)** As A, but the response variable is the fraction of PQQ DH transcripts present that contain the Ln-binding motif (fLn_T_). Trendlines are logistic regression models, and p values correspond to the x-axis coefficient.

**Fig 5.**
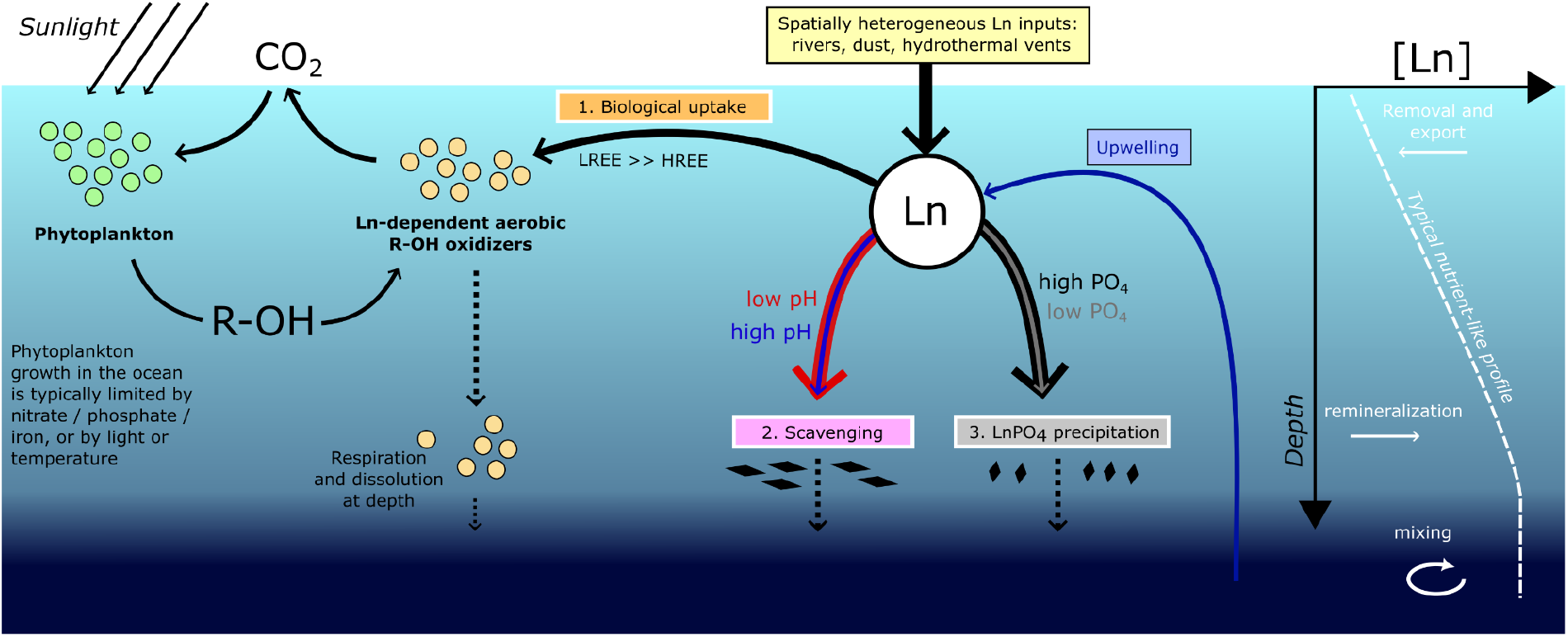
Schematic for the proposed coupling between the carbon cycle and Lanthanide cycling.

Processes 1 and 2a will increase in regions where production and export of organic matter is high. However, these effects may be counteracted by the replenishment of Ln to the surface ocean, which likely covaries with the replenishment of whichever nutrient limits phytoplankton growth rate (e.g. wind-born Fe and Ln from dust in the southern ocean^43,44^. The efficiency of adsorption reactions (2a and 2b) depends on the concentration of the free Ln^3+^ ion, which increases with decreasing pH, as a higher fraction of total Ln is in the form Ln^3+^ due to the decreased abundance of the dominant inorganic ligand, CO_3_^2-^ (Fig. S10A). This is supported by a strong positive relationship between pH and fLn_T_ (Fig. 4).

LnPO_4_ is a highly insoluble salt (K_sp_ ∼ 10^-26 -^ 10^-25^ ^45^). Rates of LnPO_4_ precipitation will be greater in waters that are more oversaturated with respect to LnPO_4_, and has been proposed to place an upper limit on the stable concentration of Ln ^13^. Process 3, therefore, depends on the solubility product [Ln^3+^][PO_4_ ^3-^]. The concentration of the PO_4_ ^3-^ ion increases with pH and total phosphate, while the free Ln^3+^ ion is a smaller fraction of total Ln at high pH^46^. The opposing effects of pH^Note 1^ on Ln^3+^ and PO_4_ ^3-^ result in an optimum relationship, with lowest concentrations of total Ln at LnPO_4_ saturation occurring at around pH 8.1, and only modest increases at lower or higher pH (Fig. S10B). This is supported by the strong negative relationships between total phosphate and fLn_T_ and fLn_T_ (Fig. 4B).

From these observations we conclude that Ln scavenging efficiency is likely most strongly controlled by pH (higher at low pH), while LaPO_4_ precipitation is most strongly controlled by phosphate (higher at high total phosphate). In the ocean, phosphate concentrations and pH negatively covary, so these removal fluxes cannot currently be decoupled. However, both processes will be more efficient where phosphate is high and pH is low such as the high-latitude HNLC regions, and less efficient where PO_4_ is low and pH is high such as the ocean gyres (Fig. 4, 5). The simplest explanation for the increased spread in fLnT with increasing phosphate concentrations is that inputs of Ln are temporally and spatially variable. At high pH and low phosphate, there is negligible depletion of surface ocean Ln, which is always available at sufficient concentrations, while at low pH and high phosphate, the steady state concentration depends on the balance between inputs and outputs.

The coupling between the marine Ln and carbon cycles likely has implications for Earth’s climate (Fig. 5). The oceanic dissolved organic carbon pool has recently been highlighted as a biogeochemically important and increasingly large reservoir of carbon ^47^, and Ln-dependent alcohol oxidation may underlie a significant leakage of carbon from this pool to the atmosphere. These interactions may drive climate feedbacks, for example, in Anthropogenic future oceans, the efficiency of Ln scavenging will likely increase following decreases in seawater pH. On longer timescales oceanic phosphate concentrations are projected to increase on millennial timescales as a result of increased weathering ^48^, which would likely cause an increase in LnPO_4_ precipitation. Both effects would drive a decrease in the surface ocean concentration of Ln, and therefore a reduction in the rate of Ln-dependent alcohol oxidation to CO_2_, manifesting as a negative feedback to increasing atmospheric pCO_2_.

### Rare earth element geochemistry

Usually referred to as the rare earth elements (REE) in the field of ocean geochemistry, the relative concentrations of Lns in seawater and in sediments have been extensively used to explore ocean circulation and hydrothermalism in the modern ocean and over geologic time ^49^. With the exception of Ce, which is oxygen sensitive, differences in the efficiency of removal fluxes between the Lns are attributed to relative stability of aqueous complexes^46^. Biological utilization of Ln in seawater, has, until very recently, been assumed to be unimportant^50^. The deep water horizon disaster in the Gulf of Mexico provided the first circumstantial evidence of biological Ln utilization. During this event, a rapid increase in water column methane concentrations drove a bloom in water column methanotrophy that was coincident with a dramatic decrease in the lightest Ln in seawater ^11^. Our results show that biological Ln utilization is not restricted to discrete events or to methanotrophy, but is ubiquitous and diverse. Microbial uptake and export thus likely imparts a previously unaccounted for fractionation of the Lns, with more efficient uptake of the light rare earths (LREE) over the heavy rare earths (HREE) which must be constrained and factored into future geochemical analyses.

Observed bacterial utilization of lanthanides has thus far only involved La to Gd, excluding Pm which does not exist in nature^24^, with a preference for the lightest Ln of the series, La, Ce and Nd. Despite being the weaker Lewis acids, and conflicting theoretical predictions of whether Ln-MDH preferentially binds the lighter^51^ or heavier^52^ of the series, selective pressures appear to favor the preferential uptake of these light Lns given their vastly higher abundances in seawater. This is likely to occur at the level of transport into the cell, perhaps in association with highly selectively binding organic ligands^53^. However, the mechanism underlying this discrimination is an open question, and an important subject for future work. The separation of lanthanides from one another remains a significant technological challenge for industry, but one that microbes appear to have already solved.

### Conclusions

Only 12 years ago the lanthanides were thought to have no role in biology, yet enzymes that depend on them are found in a fifth of all microbial genomes in the surface ocean. The dominance of the marginally more efficient Ln-PQQ-dependent enzymes over their Ca-dependent equivalents, despite the vastly higher abundance of Ca of Ln in seawater, suggests that efficiency confers a far greater advantage than the availability of their metal cofactor. Ln-dependent metabolism is an unappreciated component of the ocean microbiome that plays a central, yet entirely overlooked, role in marine carbon biogeochemistry. Uncovering the details of these biogeochemical feedbacks and implications for climatic regulation is an important aim for future research.

**Note1**: Ocean pH is buffered by carbonic acid, and changes in pH in the ocean are generally driven by changes in dissolved inorganic carbon with smaller changes in alkalinity. Discussions of the effect of changes in pH on Ln speciation is therefore calculated at constant alkalinity ^54^.

## Acknowledgments

HM acknowledges funding from the University of Melbourne. Helpful comments were provided by Max Lloyd and Tom Browning on earlier iterations of the manuscript.

## Author contributions

MV performed the bioinformatics analysis with input from JB. HM analyzed data, acquired funding and supervised the work. MV and HM conceived the study, visualized the results and wrote the first draft of the manuscript. MV, JB and HM discussed the results and edited the manuscript.

