## Supplementary Materials for "Extensive and diverse lanthanide-dependent metabolism in the ocean"

Marcos Y. Voutsinos *et al.*

**The PDF file includes:**

Materials and Methods
Supplementary Text
Figs. S1 to S10

### Materials and Methods

#### Pyrroloquinoline-quinone dehydrogenase identification and classification

We investigated the occurrence of pyrroloquinoline-quinone dehydrogenase (PQQ-DH) enzymes in the ocean by analysing several global datasets: The Tara Oceans metagenomic and metatranscriptomic Ocean Microbial Reference Gene Catalog v2.0 (OM-RGCv2), which includes includes 83 surface ocean (SO), 53 deep chlorophyll maximum (DCM), and 38 meso-pelagic zone (MES) metagenomic samples, and 103 SO, 49 DCM and 26 MES transcriptomic samples from the 0.22-3 µm size fraction [1] (Figures 1, 3 and 4 main text); and the set of 1888 Bacterial and Archaeal metagenome assembled genomes (BacArcMAGs), which spans all six size fractions of the original Tara oceans metagenomic database [2] (Figure 2 main text).

For PQQ-DH identification and classification of families we constructed a phylogenetic tree to discriminate homologous, but functionally distinct proteins that can not be identified by hidden Markov model (HMM) search alone. The PQQ-DH sequences were identified in metagenomes using a custom HMM for PQQ-binding dehydrogenases taken from [3]. Across OM-RGCv2, we identified 11,656 PQQ-DH proteins. Proteins greater than 300 amino acids in length were retained and dereplicated at 95% identity using CD-HIT v4.6 [4] resulting in 6,886 PQQ-DH sequences. These sequences were concatenated with a reference set [3,5] and aligned using FAMSA [6]. The gaps were then removed from the alignments using trimAl [7] with the parameter -gt 0.1. A phylogenetic tree was constructed using FastTree [8] and sequences were manually classified based on their relationship with the PQQ-DH reference sequences. All phylogenetic trees were constructed using the interactive tree of life (iTOL) [9]. The multiple sequence alignment was used to classify the metal binding domains by manually identifying the amino acid motifs 'D-x-A' and 'D-x-D' which represent $Ca^{2+}$ and lanthanide metal coordination respectively (**Supplementary Fig 1**). The above method was repeated for the PQQ-DH sequences derived from the dereplicated BacArcMAGs set resulting in 1,956 PQQ-DH sequences.

#### Biogeography of PQQ-DH genes and transcripts in the global ocean

The abundance of PQQ-DH genes from the global ocean were estimated by analysing the OM-RGC-v2 [1] and BAC_ARC_MAGs dataset [10]. The analysis was conducted using the Ocean Gene Atlas platform [11] with an expected threshold of 1E-10. The abundances were calculated as percent of total coverage using the reads per kilobase per million mapped (RPKM) method. The abundances of PQQ-DH transcripts were also estimated using the above methods. Only sequences with a complete cofactor binding domain were included in this analysis (calcium, lanthanide or unknown). Sequences with an incomplete cofactor binding domain (**Supplementary Fig 1, Data File S1)** were excluded.

### Genome phylogenetic classification

To taxonomically classify the microorganisms represented by the BacArcMAGs we used the combination of a concatenated ribosomal protein tree and an rpS3 protein tree. For the ribosomal protein tree, we searched each genome for 16 ribosomal proteins (RP16) using GOOSOS.py (https://github.com/jwestrob/GOOSOS). The following HMMs were used: Ribosomal_L2 (K02886), Ribosomal_L3 (K02906), Ribosomal_L4 (K02926), Ribosomal_L5 (K02931), Ribosomal_L6 (K02933), Ribosomal_L14 (K02874), Ribosomal_L15 (K02876), Ribosomal_L16 (K02878), Ribosomal_L18 (K02881), Ribosomal_L22 (K02890), Ribosomal_L24 (K02895), Ribosomal_S3 (K02982), Ribosomal_S8 (K02994), Ribosomal_S10 (K02946), Ribosomal_S17 (K02961), and Ribosomal_S19 (K02965). Ribosomal S10 model PF00338 was also used for the identification of Chloroflexi. A total of 1,669 genomes containing at least 8 ribosomal proteins were included. The ribosomal protein sequences were then individually aligned using FAMSA and concatenated using the concatenate_and_align.py script from GOOSOS (https://github.com/jwestrob/GOOSOS/blob/master/Concatenate_And_Align.py). The resulting alignments were stripped of columns containing 90% gap positions using Trimal with the parameter -gt 0.1. A phylogenetic tree was constructed using IQ-TREE [12] and the following settings: iqtree -bb 1000 -nt AUTO -ntmax 48 -mset LG+FO+R. Genomes were then classified at the phylum level using GTDB-TK [13]. Genomes with a completeness below 70% and above 10% contamination were excluded from the analysis.

### Biosynthetic gene cluster (BGC) and putative lanthanophore prediction

To identify biosynthetic gene clusters (BGCs), antiSMASH 6.0 [14] was run on the BacArcMAGs set using default parameters. Antismash identifies and annotates secondary/specialized metabolite biosynthesis gene clusters in bacterial genomes. Biosynthetic gene clusters retrieved from the genomes were dereplicated using CD-HIT at 95%. Only BGCs on contigs greater than 10 kb were included in the analysis. The antiSMASH tool only classifies BGCs as siderophores when they contain IucA/IucC genes which are specific for aerobactin and aerobactin-like siderophores. To predict the occurrence of siderophores outside of aerobactin we ran two Pfams on the BGCs pfam_transporter20.hmm and all_sbp.hmm to identify the BGCs that contain the transporters: FecCD, Peripla_BP_2, and TonB_dep_Rec. Previous work has shown these transporters are predictive of siderophore activity [15]. Putative lanthanophores were cautiously classified as such when biosynthetic gene clusters with predicted siderophore activity were detected in genomes containing lanthanide dependent enzymes. Geneious was used to visualize the biosynthetic gene clusters [16].

### Phylogenetic classification of Clade X

To identify the dominant Ln-dependent organisms a phylogenetic tree containing the most highly expressed *xoxF* genes from the metatranscriptomes with all *xoxF* genes identified in the MAGs was created (Fig. S6). Three *Alphaproteobacteria* genomes (TARA_PSW_86_MAG_00242, TARA_MED_95_MAG_00188, TARA_MED_95_MAG_00017), previously classified as Rickettsiales using GTDB-Tk and CheckM [2], contained the most highly expressed *xoxF5* sequences. They did not contain 16S sequences so they were classified based on the phylogenetic placement in a ribosomal protein tree with reference genomes taken from Luo 2015[17], Delmont et al 2022 [2] and Schön et al 2022[18]. Each genome was searched for 16 syntenic ribosomal marker proteins (rp16) using GOOSOS.py (https://github.com/jwestrob/GOOSOS). Genomes that contained at least 8 of 16 rp16 genes were retained and individually aligned using FAMSA and concatenated using the concatenate_and_align.py script from GOOSOS (https://github.com/jwestrob/GOOSOS/blob/master/Concatenate_And_Align.py). The multiple alignment was trimmed for phylogenetically informative regions using BMGE ( -m BLOSUM30) [19] and a maximum-likelihood tree was inferred using IQ-TREE [12] and the following parameters: -bb 1000 -st AA -m MFP.

**Methane monooxygenase phylogenetic analysis**

The presence of aerobic methanotrophy is typically investigated using the *pmoA* gene, which encodes subunit A of particulate methane monooxygenase (pMMO) and *mmoX*, which encodes subunit A of soluble methane monooxygenase (sMMO). The following HMMs were used to search the metagenomes: K10944 (pmoA) and K16157 (mmoX). Two phylogenetic trees were made to confirm the retrieved protein sequences. For each tree the sequences were concatenated with references taken from Singleton et al. [20] and individually aligned using FAMSA [6] with default settings. The gaps were then removed from the alignment using trimAl [7] with the parameter -gt 0.1. An approximately maximum-likelihood tree was constructed using FastTree [8] and the WAG + GAMMA parameters and sequences were manually classified based on their relationship with the MMO reference sequences. Both phylogenetic trees were constructed and annotated using iTOL[9].

**Nicotinamide adenine dinucleotide methanol dehydrogenase identification and classification**

We investigated the occurrence of nicotinamide methanol dehydrogenase (NAD MDH) genes in the global ocean by analyzing the metagenomic and metatranscriptomic Ocean Microbial Reference Gene Catalog v2.0 (OM-RGCv2) (1).

For NAD MDH identification we constructed a phylogenetic tree to discriminate homologous, but functionally distinct proteins that cannot be identified by HMM search alone. The NAD MDH sequences were identified using K00093.HMM. Across OM-RGCv2, we identified 9,043 NAD MDH proteins. Protein sequences with lengths between 320 and 420 amino acids were retained and dereplicated at 95% identity using CD-HIT v4.6 (4) resulting in 3,766 NAD MDH sequences. The sequences were concatenated with a set of experimentally confirmed NADH MDH sequences taken from [21] and aligned with FAMSA. The gaps were then removed from the alignments using trimAl (7) with the parameter -gt 0.1. The sequences were manually inspected and those that contained the NAD and metal binding domains [21] were retained, resulting in a set of 2,602 NAD MDH homologs. A phylogenetic tree was constructed using FasTree [8] and sequences were manually classified based on their relationship with the NAD MDH reference sequences. For the full phylogenetic tree see Data File S3.

**Oxygen dependent methanol dehydrogenase identification and classification**

We investigated the occurrence of oxygen dependent methanol dehydrogenase ($O_2$ MDH) genes in the global ocean by analyzing the metagenomic and metatranscriptomic Marine Atlas of Tara Oceans Unigenes (MATOU) collected during the Tara Oceans expedition [22]. This dataset was constructed to target the eukaryotic fractions of the metagenomes, including enrichment for eukaryotes and excluding prokaryotes. We present a direct comparison between $O_2$ MDH from this dataset and prokaryotic MDHs from the metagenomes / metatranscriptomes which targets free-living microbes (Fig. S8). This comparison therefore excludes potential prokaryotes in size fractions greater than 3 μm, which would only occur in association with larger particulates or in the guts of eukaryotes.

For $O_2$ MDH identification we constructed a phylogenetic tree to discriminate homologous, but functionally distinct proteins that can not be identified by only HMM search. The $O_2$ MDH sequences were identified using the K17066.HMM profile. We identified 5,491 NAD MDH proteins. Protein sequences with lengths between 190 and 700 amino acids were retained and dereplicated at 95% identity using CD-HIT v4.6 (4) resulting in 3,197 NAD MDH sequences. The sequences were concatenated with a set of $O_2$ MDH sequences taken from [23] and aligned with FAMSA. The gaps were then removed from the alignments using trimAl (7) with the parameter -gt 0.1. A phylogenetic tree was constructed using FasTree [8] and sequences were manually classified based on their relationship with the $O_2$ MDH reference sequences. For the full phylogenetic tree see Data File S4.

**Biogeochemistry**

Phosphate concentrations in Fig 4 and 5 are taken from a well established interpolated dataset of monthly average phosphate (WOA2018, NOAA), and pH was calculated from temperature, salinity, total alkalinity and DIC taken from the CO2OceanSODA-ETHZ data set [24], for the period Jan 2018 to December 2018, at 1 degree spatial resolution. To compare phosphate concentrations and pH with the metagenomic / metatranscriptomic sequence data we took the coordinates and date of the site of collection, and averaged annual averages in ocean chemistry from grid squares within a 1 degree radius of the collection site. Correlations between the fraction of DH genes that contained the Ln-binding motif (fLnG) and between the fraction of DH transcripts that contained the Ln-binding motif (fLnT) and average annual total phosphate concentrations were quantified using logistic regression analysis.

The effect of changes in pH on the solubility of $LnPO_4$ were explored at constant 15°C, and 1 bar pressure. The aqueous chemical model was developed using the python implementation of Reaktoro (reaktoro.org), using the phreeqc llnl database (thermodynamic data compiled by the Lawrence Livermore National Laboratory), and HFK activity model. A solution consisting of elements H, O, Na, Cl, C, Mg, K, S, Ca, La and P with typical seawater concentrations of ionic species: 0.468 mol / kg Na+ , 0.546 mol / kg Cl-, 0.0533 mol / kg Mg+2, 0.0281 mol / kg SO4-2, 0.0023 mol / kg HCO3-, 0.0104 mol / kg Ca+2, 0.00997 mol / kg K, 20e-12 mol / kg La and 1e-6 mol / kg PO4-3 was speciated at a temperature of 15 degrees C and 1.0 bar pressure. $LaPO_4$ was found to be the only La bearing mineral phase occurring in non-negligible amounts. Simulations involved changing $CO_2$ at constant alkalinity. Increases in $CO_2$ leads to acidification and carbonation of seawater, both of which impact Ln speciation. As pH of the water decreases, the fraction of P in the form of free $PO_4^{3-}$ and the fraction of DIC in the form of $CO_3^{2-}$ decrease. A decrease in $PO_4^{3-}$ decreases the saturation state of $LaPO_4$, however a decrease in $CO_3^{2-}$ decreases the degree of $CO_3^{2-}$ complexation of La, and therefore the fraction of La in the form $La^{3+}$ increases (Fig. S10A). These effects are counteractive for solubility of $LaPO_4$ with a minimum at around pH 8.1, but very little change with changing pH for a realistic range of ocean conditions (Fig. S10B). We conclude that the effect of pH change at constant alkalinity will likely have a significant effect on adsorptive scavenging (which depends on the $La^{3+}$ ion) but will have a negligible effect on $LaPO_4$ precipitation.

**Ordinations**

The metagenomic and transcriptomic datasets of PQQ-DHs were explored in the context of their chemical metadata. From these metadata we chose only those variables with high quality and relatively complete datasets (which did not include quality carbonate chemistry). These variables were: Temperature, Salinity, Chlorophyll A concentrations, depth, Oxygen concentration and total Phosphate. We excluded Nitrate from the analysis owing to its high collinearity with phosphate. Figure S4 presents the output from an unconstrained correspondence analysis, which was chosen to emphasize relative differences in gene abundance and expression between sites (as opposed to an unscaled analyses which is dominated by the most abundant genes / transcripts) and to avoid issues associated with non-linearity in the relationship with explanatory variables.

**Supplementary text**

Of the 33 phyla represented by the 1,888 MAGs, 14 phyla and 374 MAGs contained lanthanide dependent PQQ-DH enzymes (Ln-PQQ-DHs). A total of 1120 nonredundant Ln-PQQ-DHs lanthanide dependent PQQ-DH were identified belonging to genomes represented by Gammaproteobacteria (n = 121), Alphaproteobacteria (n = 91), Verrucomicrobia (n = 25), Bacteroidota (n = 23), Myxococcota (n = 10), Gemmatimonadetes (n = 5), Acidobacteria (n = 3), Chloroflexota (n = 3), Latescibacterota (n = 3), Planctomycetes (n = 3), Bdellovibrionota (n = 1), Campylobacterota (n = 1), Marinisomatota (n = 1), and UBP10 (n = 1). In the 1,888 MAGs, 588 genomes (31%) contained PQQ-ADH enzymes, 375 (19.8%) contained La-dep PQQ-ADH, 314 (16.6%) contained Ca-dep PQQ-ADH, and 252 (13.3%) contained unknown PQQ-ADH. And of the 1,888 genomes 168 (8.9%) contained exclusively La-dep PQQ-ADH enzymes, 128 (6.8%) contained exclusively Ca-dep PQQ-ADH, and 112 (5.9%) contained exclusively unknown PQQ-ADH enzymes.

Each of the four genomes representing Enterobacterales of Gammaproteobacteria (TARA_ION_45_MAG_00093), Rhodobacterales (TARA_PON_109_MAG_00100), Rhodospiralles (TARA_IOS_50_MAG_00003) and Tistrellales (TARA_PON_109_MAG_00031) of Alphaproteobacteria contained lanthanide-dependent enzymes and biosynthetic gene clusters containing TonB-dependent receptor, FecCD transmembrane protein (Type II ABC importer) and Peripla_BP_2 (substrate binding domain). Individually, these transporters are predictive of metallophore activity therefore these biosynthetic gene clusters are of particular interest as potential lanthanophores.

**Figures**

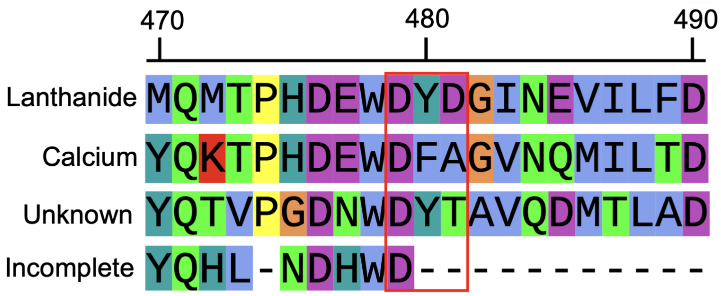


**Figure S1**: Multiple alignment of the cofactor binding domain used to classify the metal dependence of pyrroloquinoline quinone dehydrogenase sequences identified in this study. $Aspartate_{479}$ is required for PQQ binding, $aspartate_{481}$ is required for lanthanide binding, and $alanine_{481}$ is required for calcium binding. Unknown sequences classified based on the absence of Aspartate or Alanine at $AA_{481}$. Sequences classified as incomplete based incomplete binding domains.

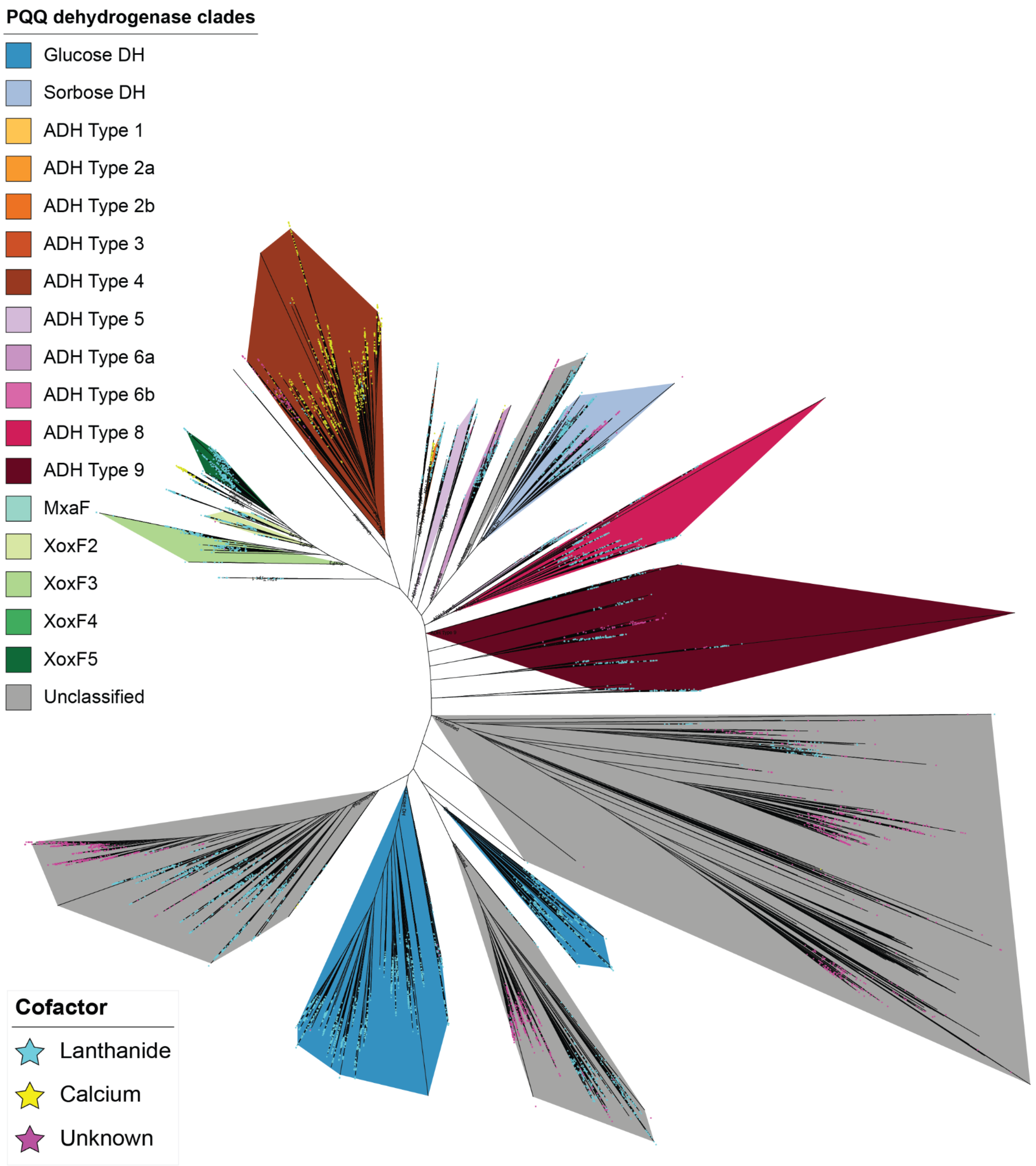


**Figure S2**: Phylogeny of the 6,886 pyrroloquinoline quinone dehydrogenase enzymes identified in the metagenomes alongside references from [5,25,26]. The metal dependence of sequences are illustrated based on the presence of stars on each leaf.

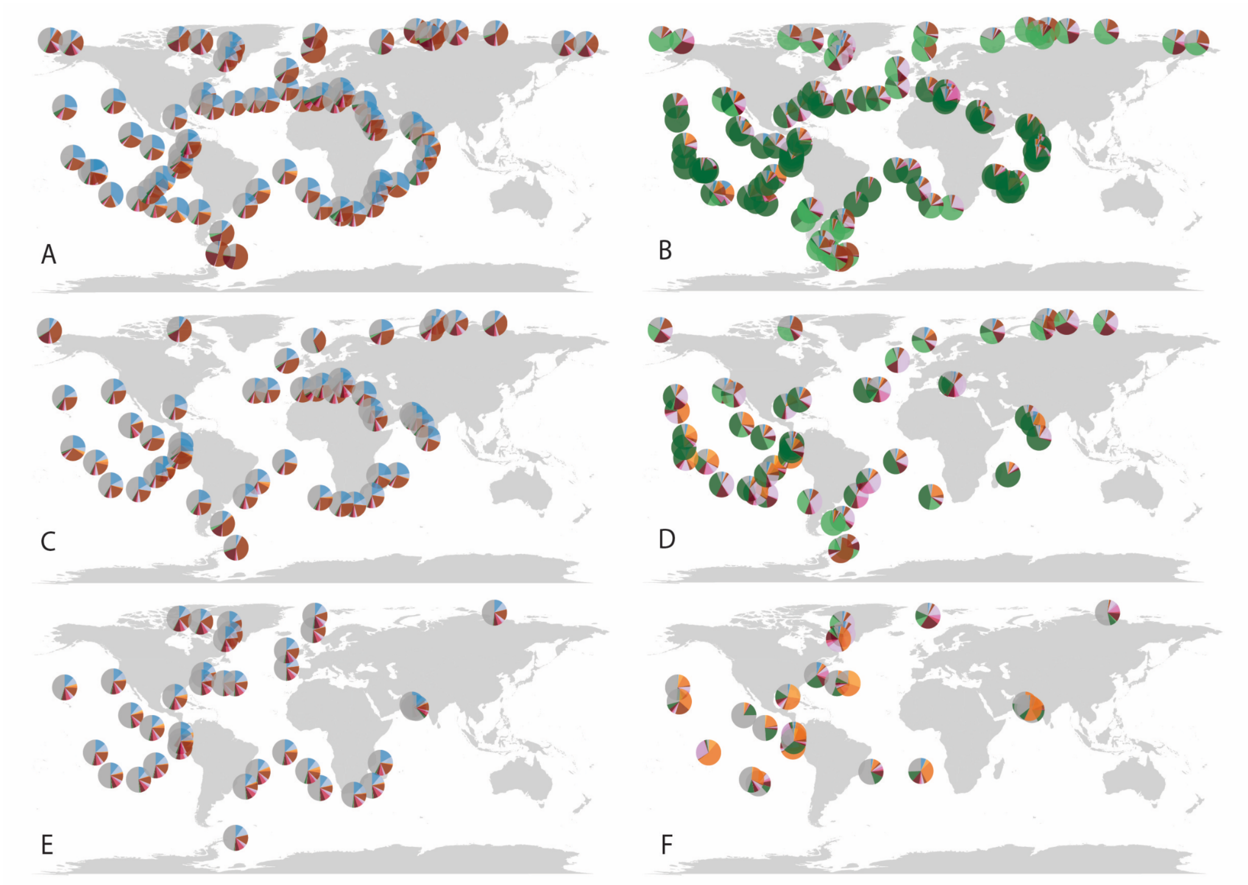


**Figure S3**: **(A)** Distribution of PQQ DH gene abundances throughout the surface ocean in community metagenomes (see figure 1C or S2 for color key). **(B)** Distribution of PQQ DH transcript abundances throughout the surface ocean in community metatrascriptomes. **(C)** Distribution of PQQ DH gene abundances throughout the deep chlorophyll maximum in community metagenomes. **(D)** Distribution of PQQ DH transcript abundances throughout the deep chlorophyll maximum in community metatranscriptomes. **(E)** Distribution of PQQ DH gene abundances throughout the mesopelagic zone in community metagenomes **(F)** Distribution of PQQ DH transcript abundances throughout the mesopelagic zone in community metatrascriptomes.

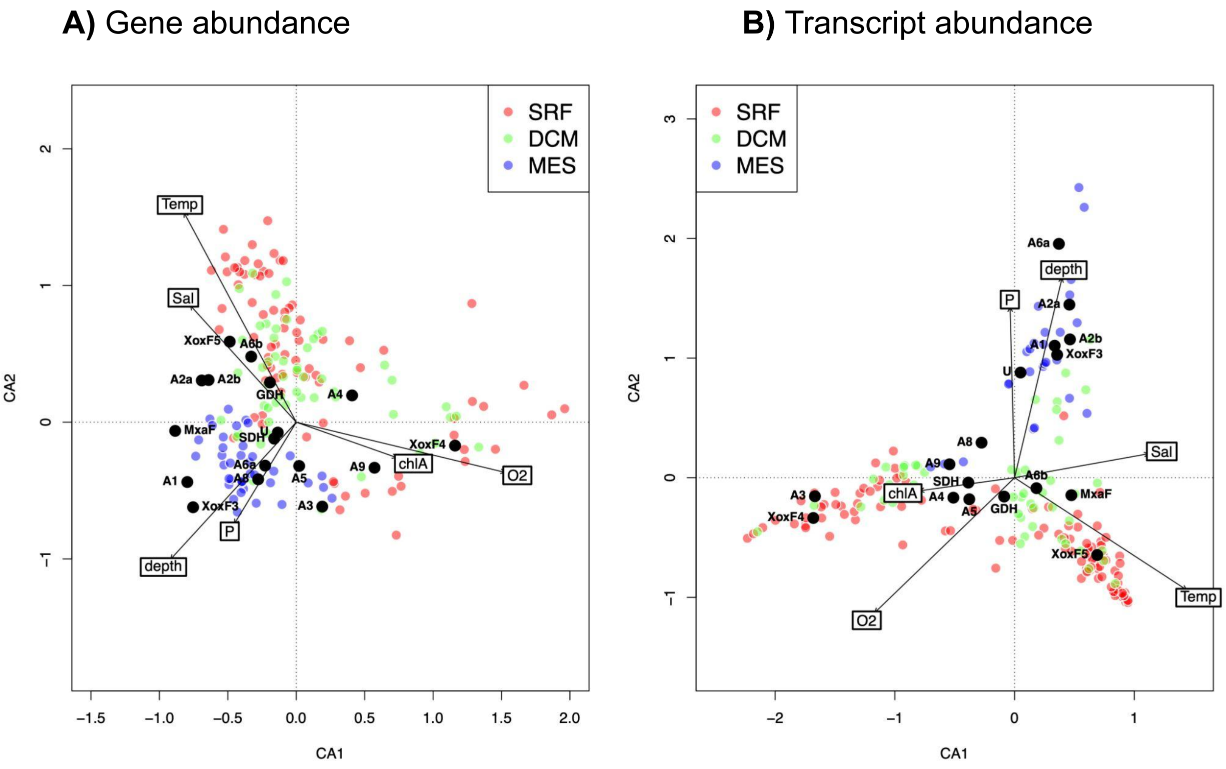


**Figure S4.** Triplots showing the results of an unconstrained correspondence analysis on A) gene abundances and B) transcript abundances, with environmental variables. The transcript abundance ordination produces 3 distinct regimes: 1. High temperature surface samples (bottom right arm), 2. Low temperature surface samples (left arm), and 3. Deep samples (upper arm). The sampled from the DCM cluster between these extremes.

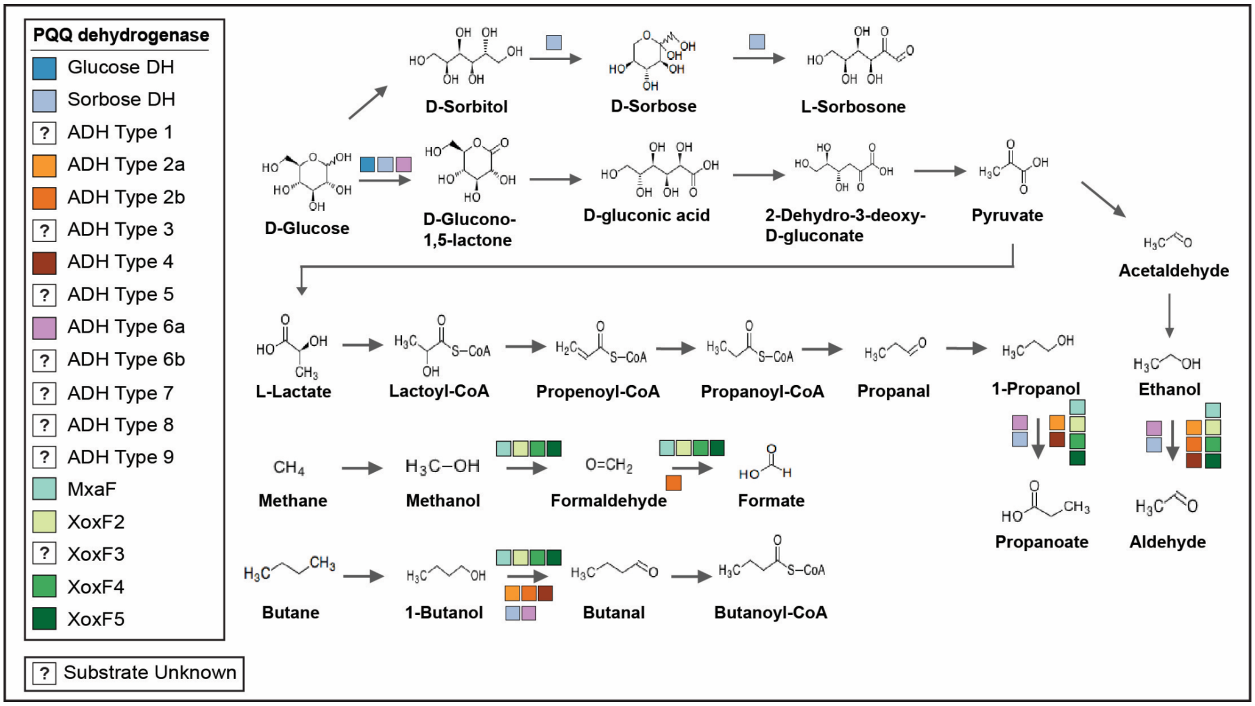


**Figure S5:** Summary of experimentally confirmed metabolic activities of pyrroloquinoline quinone dehydrogenases that were identified in this study. See data file S2 for references.

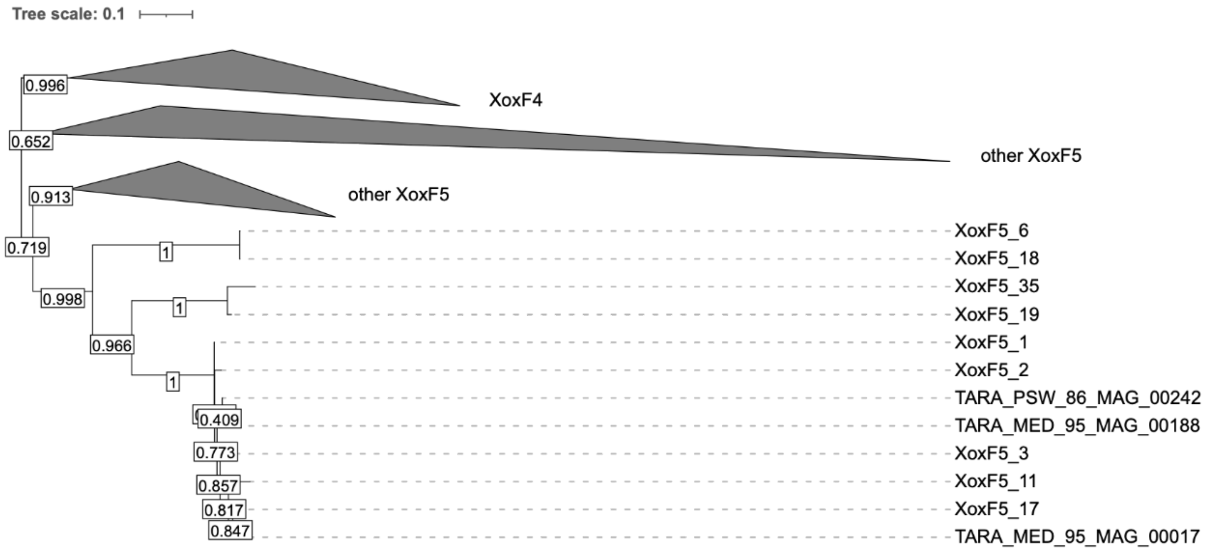


**Figure S6**: Phylogeny of the most highly expressed XoxF sequences from the metatranscriptomes and XoxF sequences from the MAGs. Metatranscriptomic XoxF5 sequence labels amended with abundance rank.

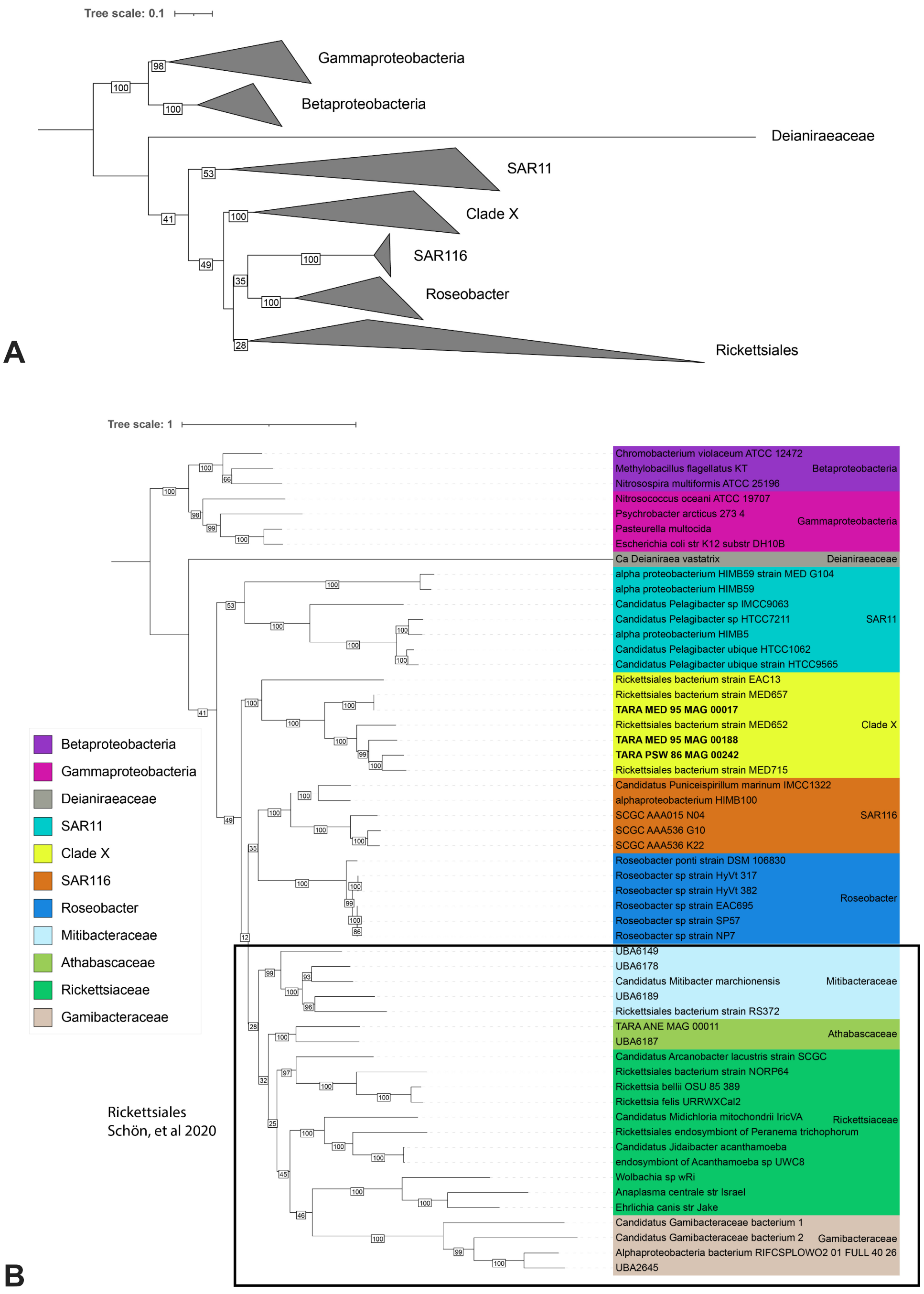


**Figure S7**: **A**) Collapsed phylogenetic tree showing the placement of Clade X with sister clades. Clades were collapsed based on a <50 bootstrap value. **B**) Expanded phylogenetic tree of the three Alphaproteobacteria genomes (Clade X) identified in Figure S6 to contain *xoxF5* sequences that group closely with the most highly expressed *xoxF5* sequences. The phylogenetic tree was constructed with a concatenated alignment of 16 ribosomal phylogenetic marker proteins (see methods). Reference genomes are taken from Luo 2015, Delmont et al 2022 and Schön et al 2022.

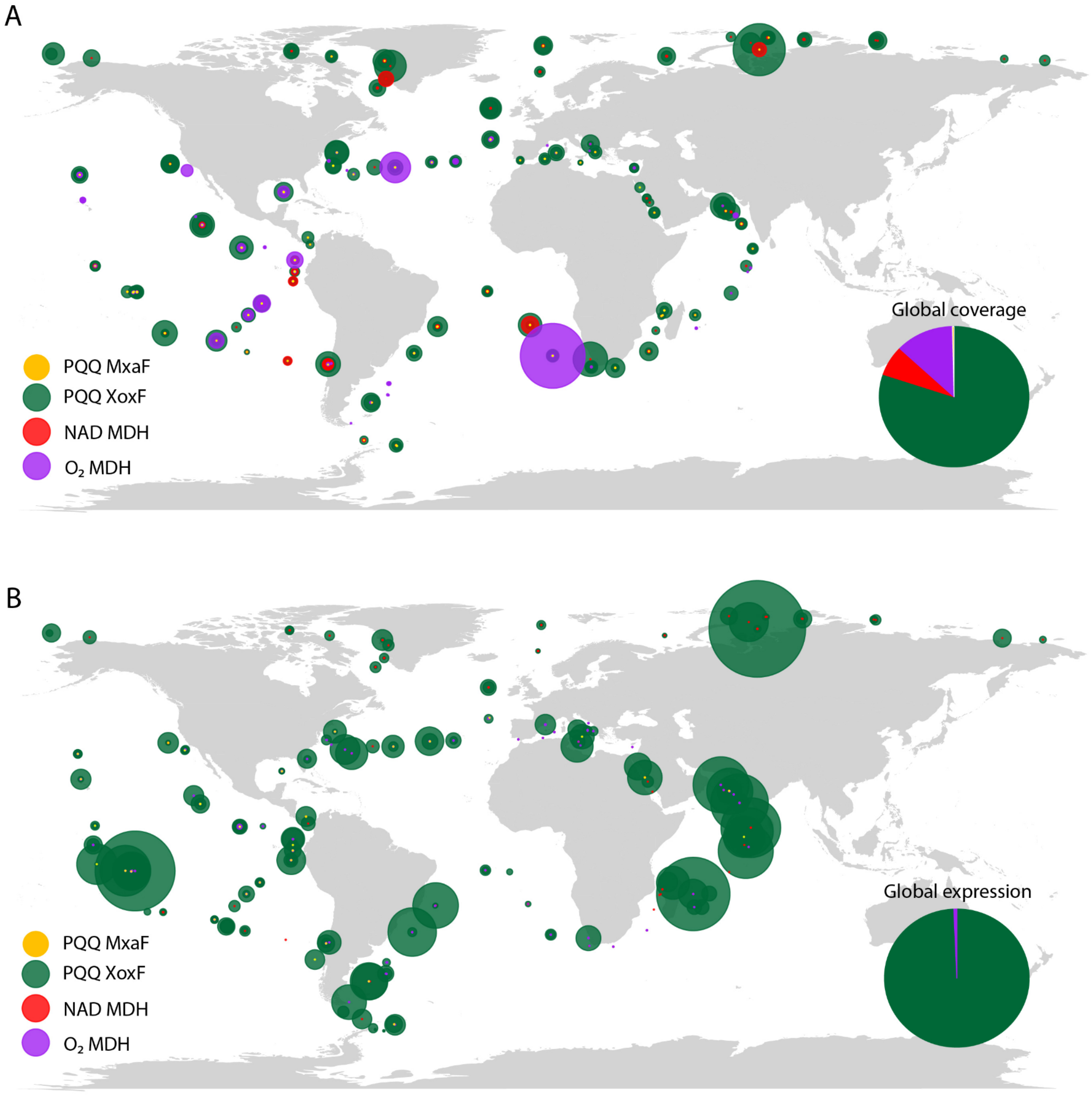


**Figure S8:** Distribution of PQQ MxaF, PQQ XoxF, NAD MDH and $O_2$ MDH **(A)** gene abundances and **(B)** transcript abundances in the surface ocean, deep chlorophyll maximum and mesopelagic. PQQ MxaF, PQQ XoxF and NAD MDH abundances were derived from the 0.22-3 µm size fraction and $O_2$ MDH abundances were derived from the 0.8-2000 µm size fractions.

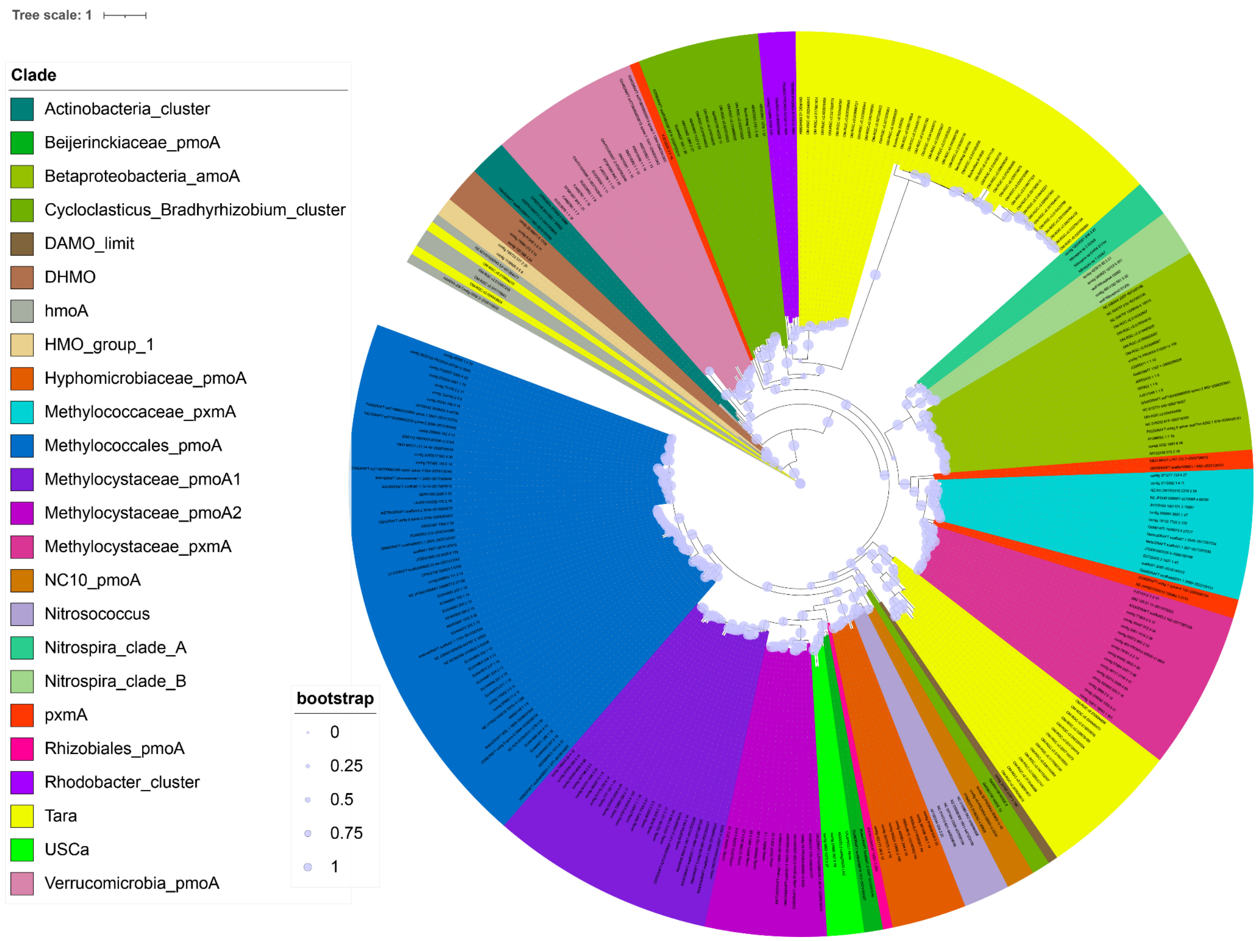


**Figure S9a:** Phylogeny of homologous PmoA proteins recovered from the metagenomes. Tara sequences analysed in this work are highlighted in yellow. Reference sequences from Singleton et al. [20].

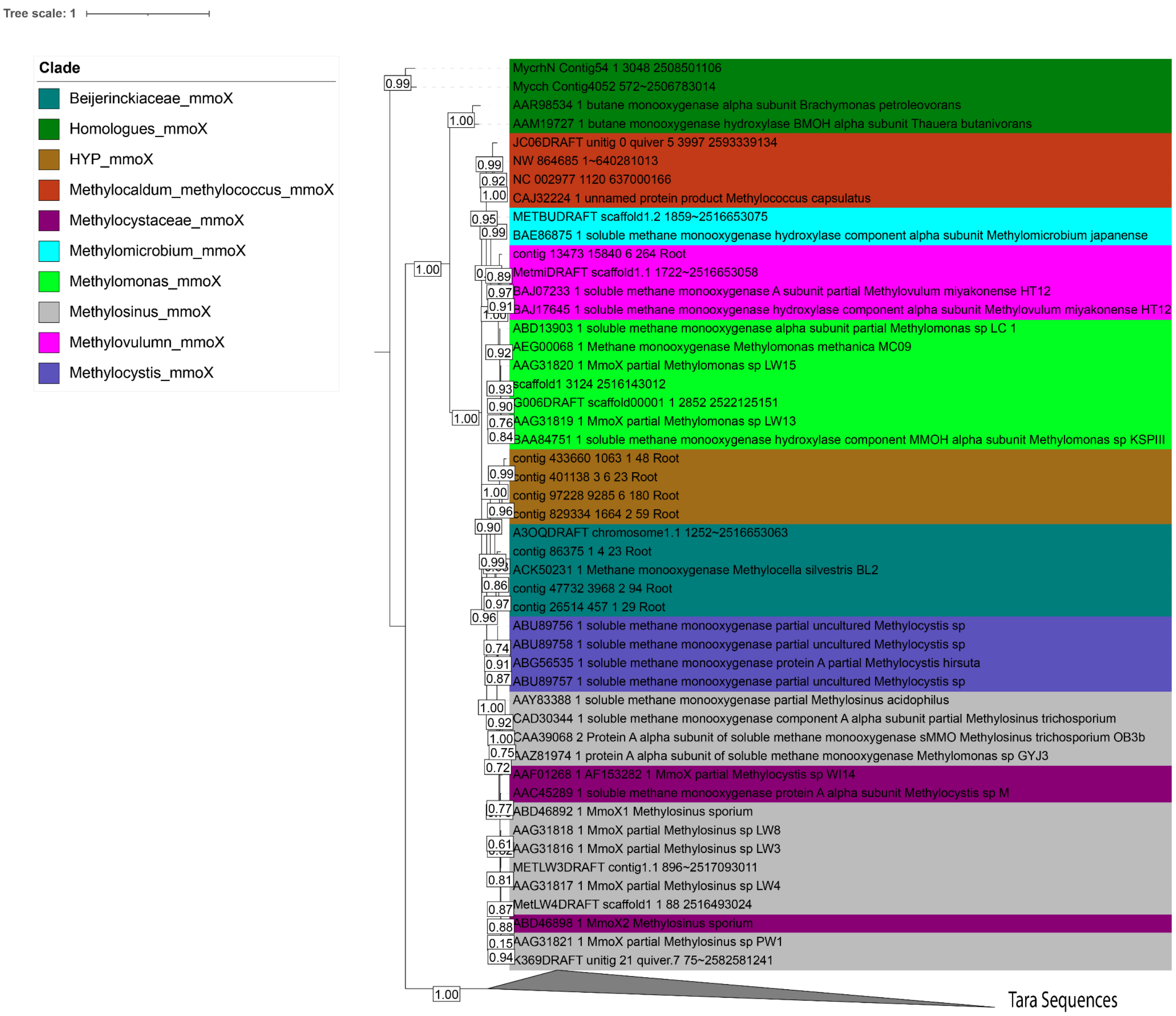


**Figure S9b:** Phylogeny of homologous MmoX proteins recovered from the metagenomes. Reference sequences from Singleton et al. [20].

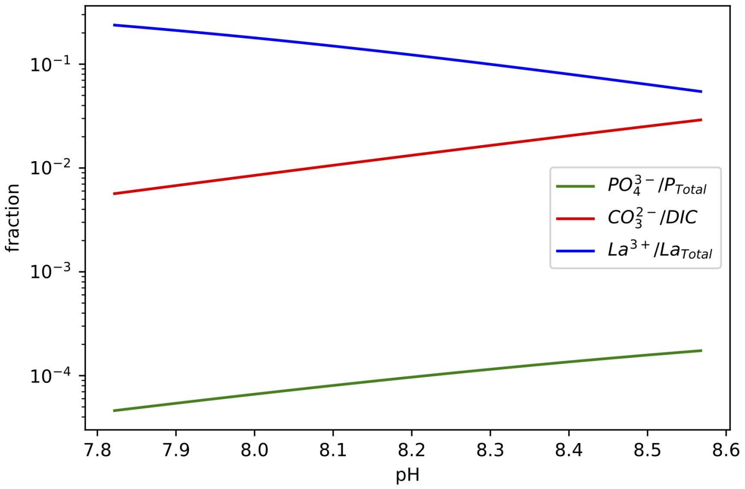


**A)**

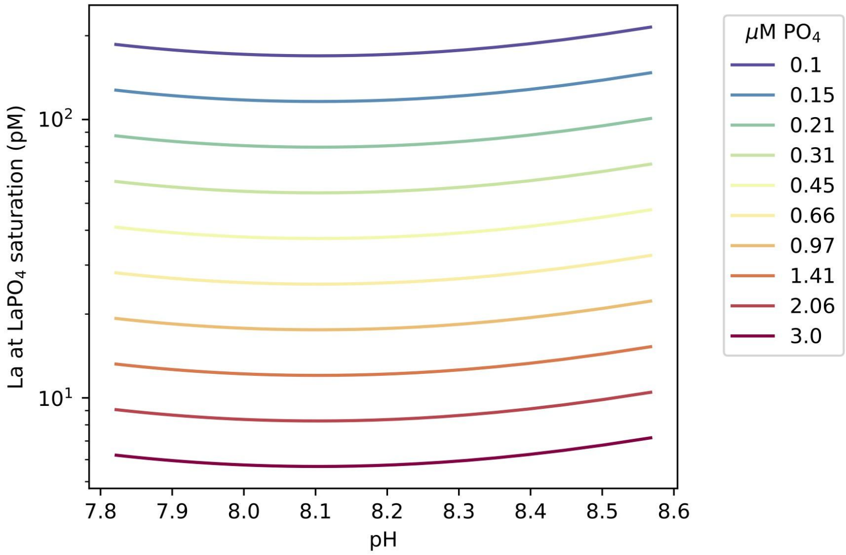


**B)**

**Figure S10: A**) Representation of P, DIC and La (as representative of Ln) speciation with pH under typical surface ocean conditions. Note $La^{3+}$ / $La_{Total}$ increases strongly with decreasing pH. **B)** Total La concentrations at $LaPO_4$ saturation. Where La concentrations are higher than this value, $LaPO_4$ will precipitate. The concentration of La at LaPO4 saturation is only weakly impacted by pH, but is strongly controlled by total phosphate concentrations. In both plots pH is varied by changing total dissolved inorganic carbon at constant alkalinity.
